# ComF is a key mediator in single-stranded DNA transport and handling during natural transformation

**DOI:** 10.1101/2021.08.04.455028

**Authors:** Prashant P. Damke, Louisa Celma, Sumedha Kondekar, Anne Marie Di Guilmi, Stéphanie Marsin, Jordane Dépagne, Xavier Veaute, Pierre Legrand, Hélène Walbott, Julien Vercruyssen, Raphaël Guérois, Sophie Quevillon-Cheruel, J. Pablo Radicella

**Affiliations:** Université Paris-Saclay, CEA, Institut de Biologie François Jacob, Stabilité Génétique Cellules Souches et Radiations, F-92260 Fontenay aux Roses, France; Université de Paris, CEA, Institut de Biologie François Jacob, Stabilité Génétique Cellules Souches et Radiations, F-92260 Fontenay aux Roses, France; Université Paris-Saclay, CEA, CNRS, Institute for Integrative Biology of the Cell (I2BC), F-91198 Gif-sur-Yvette, France; Université Paris-Saclay and Université de Paris, CEA, INSERM, Institut de Biologie François Jacob, Stabilité Génétique Cellules Souches et Radiations, F-92260 Fontenay aux Roses, France; Synchrotron SOLEIL, L’Orme des Merisiers, F-91192 Gif-sur-Yvette, France

## Abstract

Natural transformation plays a major role in the spreading of antibiotic resistances and virulence factors. Whilst bacterial species display specificities in the molecular machineries allowing transforming DNA capture and integration into their genome, the ComF(C) protein is essential for natural transformation in all Gram-positive and - negative species studied. Despite this, its role remains largely unknown. Here, we show that *Helicobacter pylori* ComF is not only involved in DNA transport through the cell membrane, but it also required for the handling of the ssDNA once it is delivered into the cytoplasm. ComF crystal structure revealed the presence of a zinc-finger motif and a putative phosphoribosyl transferase domain, both necessary for its *in vivo* activity. ComF is a membrane-associated protein with affinity for single-stranded DNA. Collectively, our results suggest that ComF provides the link between the transport of the transforming DNA into the cytoplasm and its handling by the recombination machinery.

## Introduction

Bacterial populations display an amazing capacity to adapt to changes in their environment. In pathogens, this is reflected in the generation of variants able to colonise new hosts, the propagation of virulence factors or the acquisition antibiotics resistance. A key mechanism in the propagation of those traits is horizontal gene transfer (HGT). There are three major mechanisms of HGT in bacteria: conjugation, phage transduction and natural transformation (NT). Although NT has been documented in at least 80 bacterial species ^1^, many aspects of the underlying mechanisms and players remain to be unveiled. Unlike the other pathways of HGT, NT only requires proteins coded by the recipient cell. It relies on the presence in naturally competent bacteria of a sophisticated apparatus capable of capturing DNA present in the environment and integrating it into their chromosome ^2^.

In both gram-positive and -negative species, NT can be divided in four distinct steps ^2^. It is initiated by the capture of exogenous double-stranded DNA (dsDNA) molecule at the surface of the cell where it binds to macromolecular complexes. During the uptake step, dsDNA is imported from the surface and into the periplasm (defined here as the compartments between the outer and inner membranes in gram-negative bacteria and between the cell wall and the membrane in gram-positive bacteria ^2^). In the periplasm, the incoming DNA is directed to the inner cell membrane for its transport into the cytoplasm as single-stranded DNA (ssDNA). There, it is handled to the recombination machinery leading eventually to its incorporation into the chromosome, the last step of the process.

At each step of NT specialised proteins are required. Some of them are common to most species studied, but others are species-specific. While type IV pseudo-pili components have been shown to mediate the binding of the DNA to the cell surface in *Streptococcus pneumoniae* ^3^, this observation cannot be generalised to all competent bacteria. In *Bacillus subtilis*, wall teichoic acids, but not the pseudo-pilus, are involved in the initial binding of the DNA ^4,5^. Actually, in most cases the molecules responsible for DNA capture are still unidentified. *Helicobacter pylori*, a species characterised for its high capacity for NT, does not harbour genes coding for type IV (pseudo-)pili. As for the uptake step, in nearly all the naturally transformable bacteria it is mediated by a type IV pseudo-pilus ^2,6^, with the exception of *H. pylori* that uses a type IV secretion system to pull the DNA into the periplasm ^7–9^. To complete this step, the conserved DNA receptor ComEA, a periplasmic or membrane-associated protein in gram-negative or gram-positive bacteria, respectively, is required. Here again, *H. pylori* constitutes an exception, since a unique protein, ComH, takes this role ^10^. The transport step is carried out by the membrane channel ComEC ^6,9,11–13^. ComEC is present and required for NT in all naturally transformable bacteria. Finally, the handing of the incoming DNA to the recombination machinery requires DprA ^14^, another transformation-specific protein with orthologues in all naturally transformable species.

Other proteins essential for NT have been identified by genetics approaches, but their role is still unknown. One of them, ComF(C), is present in all naturally competent bacteria. The requirement of this protein for transformation was discovered almost 30 years ago ^15,16^. The *B. subtilis* ComF locus consists of an operon harbouring three open reading frames coding for the proteins ComFA, ComFB and ComFC. All three are required for competence ^16,17^. ComFA, which appears to be present in all naturally transformable Gram-positive bacteria but has not been identified in Gram-negative species, is a DNA-dependent ATPase ^18–20^. A regulatory function has been proposed for ComFB ^17^. In the case of ComFC, its function remains unknown despite being the only one of the three for which orthologues have been identified and described as essential for competence in all naturally transformable species studied.

Here, we characterised the ComF(C), herein ComF, from *H. pylori* and investigated its role in NT. Through the determination of its 3D structure we found that ComF harbours phosphoribosyl transferase (PRT) and Zn-finger domains, both essential for transformation. We show that in the absence of ComF, not only the transport of the transforming DNA (tDNA) into the cytoplasm is blocked but also its integration into the bacterial chromosome is impaired when the DNA is directly delivered to the cytoplasm. These phenotypes, together with the observations of ComF association with the inner cell membrane and its capacity to bind ssDNA suggest a model in which ComF role provides a link between the transport and recombination steps during NT.

## Results and discussion

### ComF participates in DNA transport through the inner membrane

Despite the critical role of natural transformation in bacterial evolution and in the propagation of virulence and antibiotic resistances, many aspects of the transforming DNA uptake and processing remain poorly understood. A notorious example is the role of the ComF protein, which, while its gene was identified almost 30 years ago as essential for competence ^15,16^ and conserved in all naturally transformable species studied so far, remains unknown. A transposon mutagenesis screen originally identified *hp1473 (comF)*, a *comFC* orthologue, as a gene essential for natural transformation in *H. pylori* ^21^. In the present study we undertook the characterisation of this protein and its function by a combination of approaches to (i) define the step(s) at which the protein acts during NT, (ii) determine its 3D structure and (iii) analyse its biochemical properties.

To confirm the effect of *comF* inactivation on *H. pylori* NT, a *hp1473* null mutant was generated by insertion of a non-polar cassette and NT frequencies were determined using genomic DNA from a streptomycin resistant strain as transforming DNA. The absence of ComF led to almost a four log decrease in the transformation efficiency when compared to the wild-type strain (Fig. 1a). This phenotype, although less severe than that induced by inactivation of *comB, comEC* or *recA*, was similar to that observed for *ΔcomH* and *ΔdprA* strains. Wild-type levels of transformation were restored by the re-expression of *comF* gene introduced with its own promoter at the *rdxA* locus (Fig. 1b), ruling out polar effects of the deletion.

**Fig. 1:**
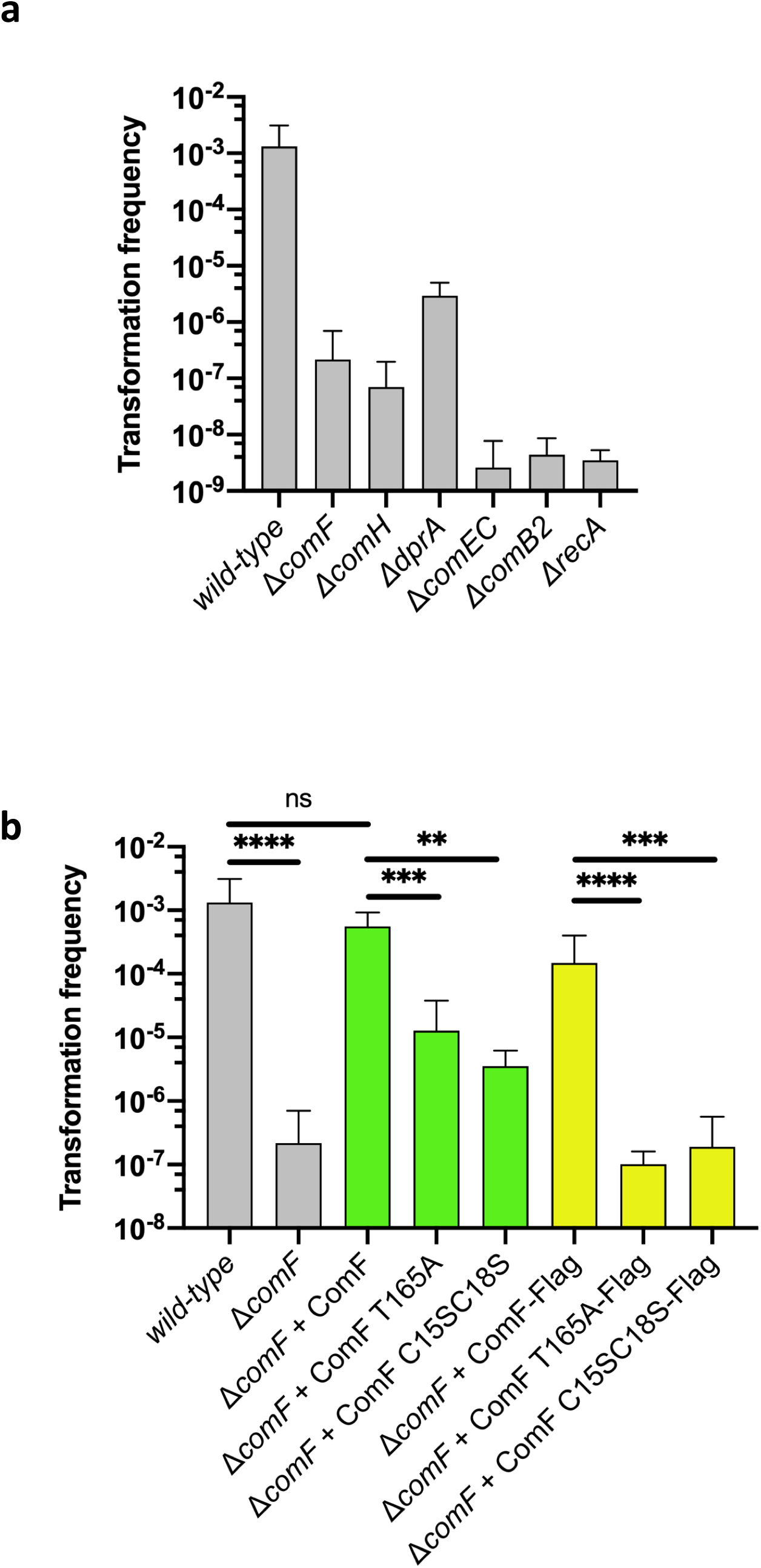
ComF is essential for genetic transformation of *H. pylori*. (**a**) Natural transformation frequencies for indicated *H. pylori* strains. (**b**) Complementation of the Δ*comF* strain. Natural transformation frequencies for indicated *H. pylori* strains were determined using isogenic streptomycin resistant total genomic DNA as donor. Bars correspond to the mean and standard deviation from at least three independent biological replicates. ns, not significant (*P* > 0.05); \*\**P* < 0.01, \*\*\**P* < 0.001 and \*\*\*\**P* < 0.0001. *P* values were calculated using the Mann–Whitney *U* test on GraphPad Prism software.

We then sought to define at which stage(s) during the transformation process ComF is required. Deletion of *comF* did not affect the uptake step as illustrated by the presence of transforming DNA foci (Fig. 2a and b). Consistently with the cytoplasmic localisation of ComF and the two-steps model for DNA uptake ^9,22^, the wild-type levels of DNA foci in *ΔcomF* strains indicated that ComF is not required for the exogenous DNA capture and uptake into the periplasm. The same conclusion was reached by monitoring by PCR the presence of tDNA in *ΔcomF* mutant strains of *V. cholerae* ^6^. However, when the persistence of the foci was monitored, we observed that in the *ΔcomF* strain foci are detected for longer times than in the wild type (Fig. 2b and c), similar to what was observed in a *comEC* mutant ^12^. This suggested that ComF is needed for an efficient transport of the incoming DNA through the inner membrane. Consistently, when the kinetics of fluorescent DNA internalisation were followed in living bacteria ^12^, we observed that in the *ΔcomF* mutant, as it is the case in the *ΔcomEC* one, the transforming DNA could not be detected as entering into the cytoplasm (Fig. 2d and Supplementary Movies 1-3). These observations are very similar to those described in *ΔcomEC* strains ^9,12^, suggesting that ComEC and ComF act at the same NT step to mediate DNA transport across the inner membrane. Taken together, these results indicate that ComF participates in the passage of the transforming DNA into the cytosol.

**Fig. 2:**
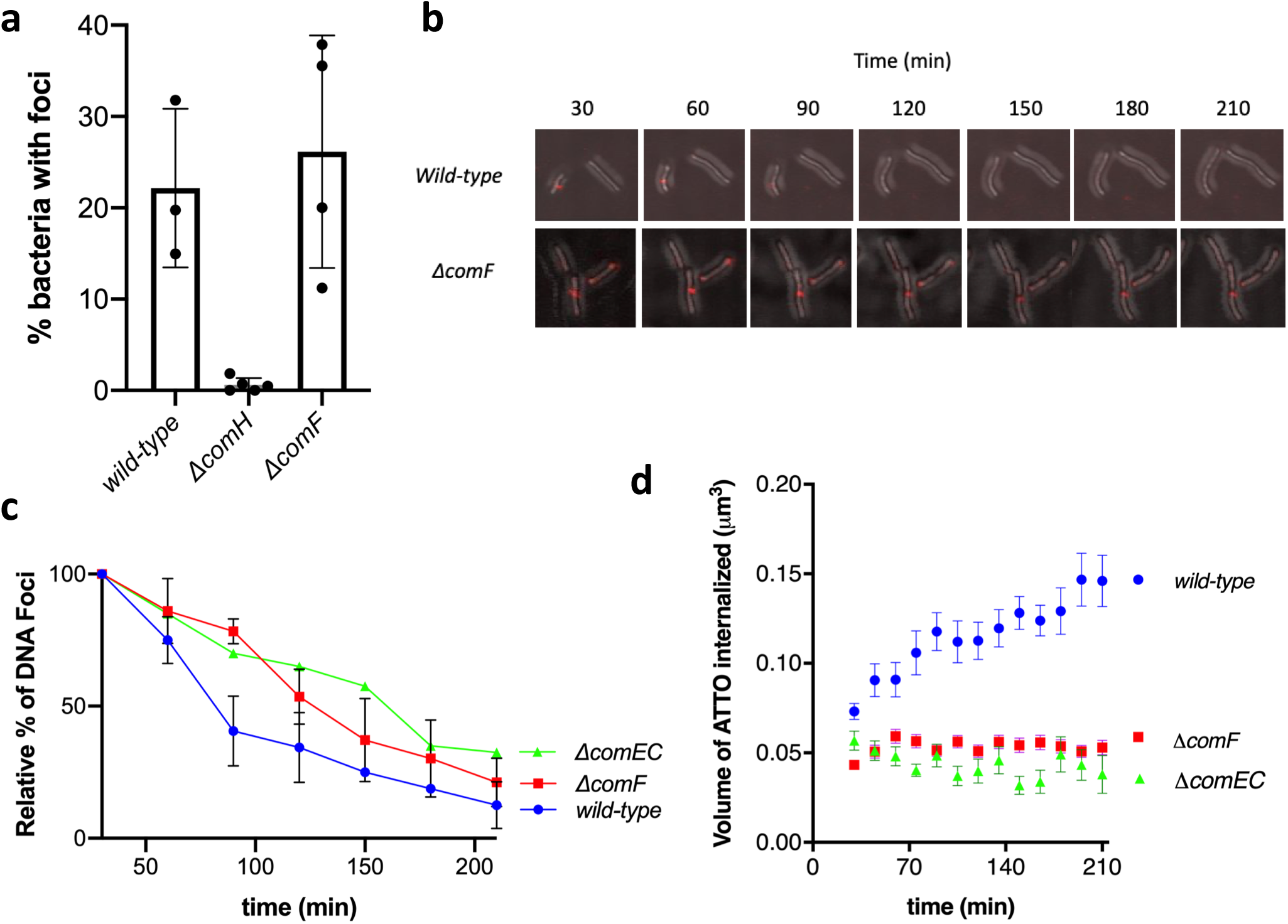
ComF supports translocation of tDNA across the cytoplasmic membrane. (**a**) Percentage of cells with fluorescent DNA foci for indicated *H. pylori* strains. Bars correspond to the average and standard deviation from at least two independent biological experiments. **(b)** Time course of DNA foci for indicated *H. pylori* strains. Z maximum projections of merged images of ATTO-550 (red channel) and differential interference contrast (DIC) are presented. **(c)** Stability of DNA foci displayed by *H. pylori* strains. Data points correspond to the mean and standard deviation from at least two independent experiments except for the *ΔcomEC* strain. **(d)** Internalization kinetics of fluorescent DNA in indicated *H. pylori* strains. GFP expressing bacteria displaying fluorescent DNA foci were followed for 3 h by confocal microscopy in live conditions (Supplementary Movies 1-3). The mean +-SEM volumes of DNA internalized were measured by 3D-analysis of individual bacterial cells for wild-type (n=26), *ΔcomEC* (n=25), *ΔcomF* (n=148) strains. At least two independent experiments were performed for each strain except for *ΔcomEC. P*-values calculated using Kruskal–Wallis statistics indicate that Δ*comEC* (*p*<0.0001), ΔdprA (*p*<0.0050), and *ΔcomF* (*p*=0.0003) curves are significantly different from the wild-type curve.

### ComF is associated with the inner membrane

A role of ComF in the tDNA translocation from the periplasm to the cytoplasm supposes a connection of the protein with the membrane. To explore such a possibility, we developed an antibody against ComF. Initially we were not able to detect the protein by immunoblot with this antibody, but by skipping the boiling step, a specific, albeit weak, signal was detectable in the wild-type strain extract (Supplementary Figure 1). We then fractionated the extracts into soluble and membrane fractions and we observed that ComF was associated with the membrane compartment (Fig. 3a). Further fractionation showed that ComF co-purified with the inner membrane fraction (Fig. 3b). To obtain a better signal, ComF fused to a FLAG tag (ComF-FLAG) was ectopically expressed from the *ureA* promoter. ComF-FLAG complementation of the *comF* deletion was less efficient, but could still support high levels of transformation (Fig. 1b). ComF-FLAG was also present in the membrane fraction, although it could also be found in the soluble fraction probably due to its overexpression (Fig. 3c).

**Fig. 3:**
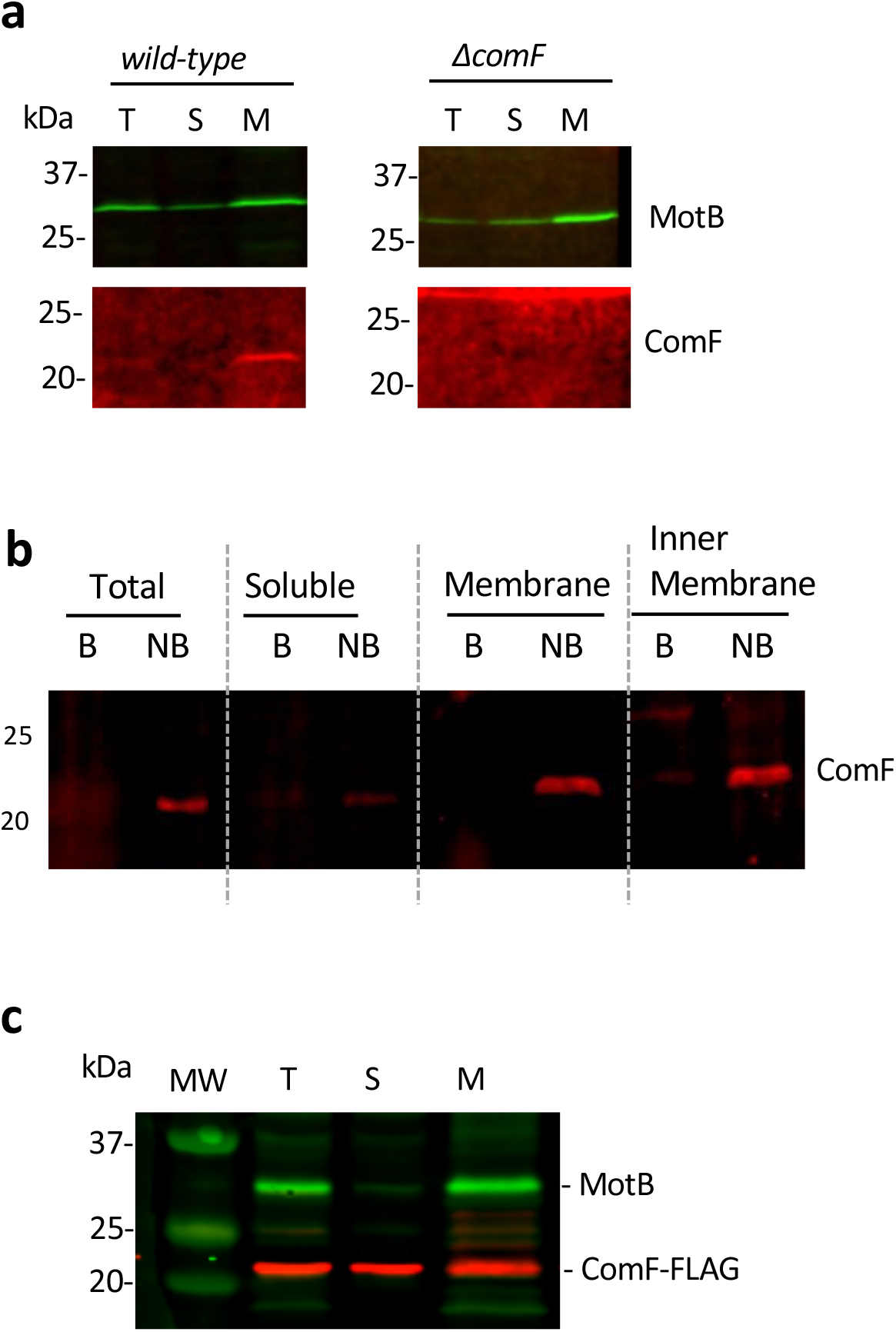
ComF is a membrane-associated protein. (**a**) and (**b**) Localisation of ComF in wild-type *H. pylori* strain. (**c**) Localisation of overexpressed ComF-FLAG. The inner membrane protein MotB was used as marker for the fractionation experiments. T: total extract, S: soluble fraction, M: membrane. B: boiled samples, NB: not boiled samples.

The association of ComF with the membrane is consistent with the role of the protein facilitating the transport of the exogenous DNA through the inner cell membrane. The link to the membrane could be direct or mediated by either another protein or the incoming DNA. A candidate for coupling ComF to the membrane in Gram-positive bacteria could be ComFA. In *B. subtilis* ComFA was shown to be a membrane protein ^19^. Furthermore, it was shown that its *S. pneumoniae* orthologue interacts with ComF(C) ^18^. However, no orthologue of ComFA has been so far found in Gram-negative naturally transformable bacteria where ComF could interact with a functional, but yet to be identified, homologue of ComFA. Alternatively, ComF targeting to the membrane could be mediated by a completely unrelated protein. Finally, although unlikely in view of the lack of a membrane anchoring domain, ComF could be itself binding to the periphery of membranes.

### ComF is required for tDNA handling within the cytoplasm

While the experiments described above demonstrated a role of ComF in the internalisation of the transforming DNA, they do not rule out its involvement in downstream steps of the natural transformation process. In order to test this possibility, the internalisation step, impaired in the *ΔcomF* mutant, needs to be bypassed. The transforming ssDNA was therefore delivered to the cytoplasm by electroporation. While ssDNA is a poor substrate for NT ^23^, electroporation with a 75-mer ssDNA carrying a streptomycin resistance marker allowed transformation of mutants deficient in the uptake (*ΔcomB2*) and internalisation (*ΔcomEC*) steps ^10^. However, after electroporatin with the same ssDNA, either much less or no streptomycin resistant transformants was observed for mutants affecting the homologous recombination process (*ΔdprA, ΔrecA*) (Fig. 4a). When a *ΔcomF* mutant was electroporated with the same ssDNA, the level of streptomycin resistant recombinants was similar to that obtained with a *ΔdprA* strain. Similar results were obtained with a longer substrate (139-mer also carrying the mutation conferring streptomycin resistance) (Supplementary Figure 2). The reduced efficiency of a *ΔcomF* mutant in transformation by electroporation with single-stranded DNA suggests that ComF is involved in NT steps downstream of the transport through the inner membrane.

**Fig. 4:**
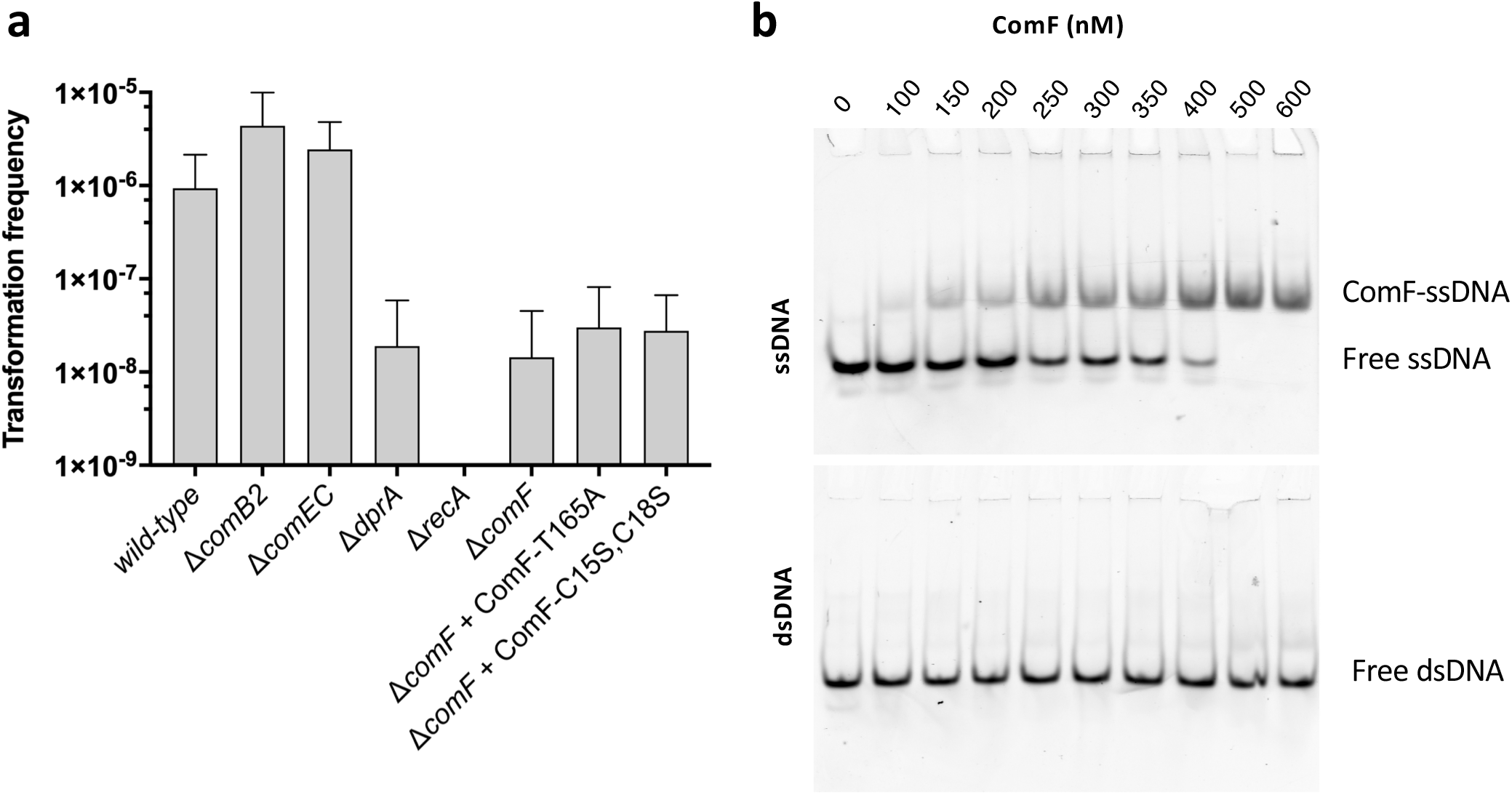
ComF binds single-stranded DNA and promotes its chromosomal integration. (**a**) Transformation frequencies after electroporation with a chemically synthesized single-stranded DNA (75 -mer) coding for streptomycin resistance as donor DNA. Bars correspond to the average and standard deviation from at least two independent biological experiments. (**b**) Selective affinity of ComF for single-stranded DNA. Indicated concentrations of His_6_-ComFC were incubated with Cy5 labelled single- or double-stranded DNA substrate. The nucleoprotein complexes were resolved by native-PAGE.

To further explore this role of ComF in the handling of the tDNA in the cytoplasm, we purified ComF and analysed its capacity to bind DNA by electrophoretic mobility shift assays (EMSA). ComF formed discrete nucleoprotein complexes with a 62-mer single-stranded DNA (ssDNA) in a concentration-dependent manner while no binding to the corresponding dsDNA was detectable (Fig. 4b). ComF bound single-stranded oligonucleotides with relatively high affinity (half-maximal binding concentration of 300 nM). This marked preference for ssDNA is consistent with the fact that during NT the incoming DNA enters the cytoplasm as ssDNA ^24^.

The failure to bypass the transformation defect of mutant *ΔcomF* by electroporation with ssDNA together with the capacity of ComF to bind ssDNA, indicate that ComF is likely to be implicated in the steps leading to the formation of the recombination substrate within the cytoplasm. ComF could, together with DprA and RecA ^25^, participate in the protection of the incoming DNA from degradation. Interestingly, ComF from *Campilobacter jejuni* and ComF(C) from *S. pneumoniae*, both required for natural transformation ^18,26^, interact with DprA ^18,27^. Since DprA plays a critical role in the loading of the recombinase to the transforming DNA ^14^, it is tempting to speculate that ComF binds the tDNA emerging from ComEC into the cytoplasm and targets it to DprA to protect it from degradation and allow further processing by the recombination machinery.

### ComF harbours zinc-finger and PRT domains

Despite its conservation amongst naturally competent bacteria (Supplementary Figure 3), no structural data on ComF(C) proteins is available. The determination of its 3D structure has been elusive. After unsuccessful crystallisation attempts with the isolated protein we generated a gene fusion between the full-length *comF* gene and an artificial αRep binder coding sequence selected from a highly diverse library of artificial repeat proteins based on thermostable HEAT-like repeats in order to help cristallisation ^28,29^. Since structural homology predictions using HHpred (toolkit.tuebingen.mpg.de/hhpred/)^30^ suggested that the C-terminal domain of ComF (residues 53 to 188) harbours a putative nucleoside binding site characteristic of phosphoribosyl transferases (PRTases) belonging to the PRT family (PurF, PDB number 6CZF-A, probability 99.15%), crystals were grown in the presence of 5-phospho-α-D-ribosyl 1-pyrophosphate (PRPP). Diffracting crystals were obtained with the purified fusion protein and the 3D structure of full-length ComF in complex with PRPP was solved at 2.56 Å resolution (Table 1, Materials and Methods) ^31^. The fusion (αRep: residues 1–229, linker: residues 230– 236, ComF: residues 237-427) is present in four copies in the asymmetric unit, organised in two domain-swapped dimers: the αRep of one fusion covers the ComF of the other one (Fig. 5a). The 3D structure obtained confirmed the presence of two predicted distinct domains in ComF.

**Table 1.**
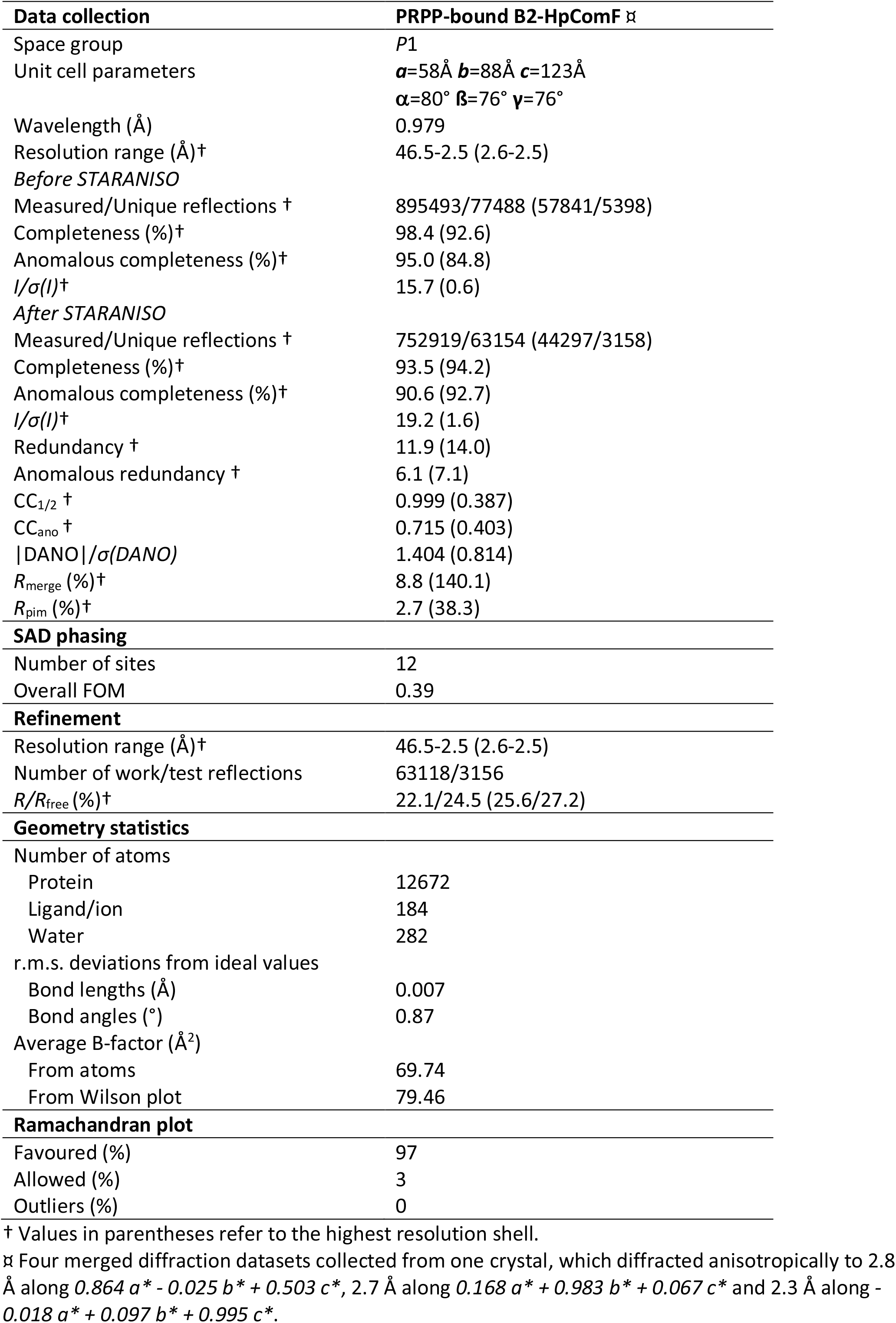
Data Collection and Structure Refinement Statistics.

**Fig. 5:**
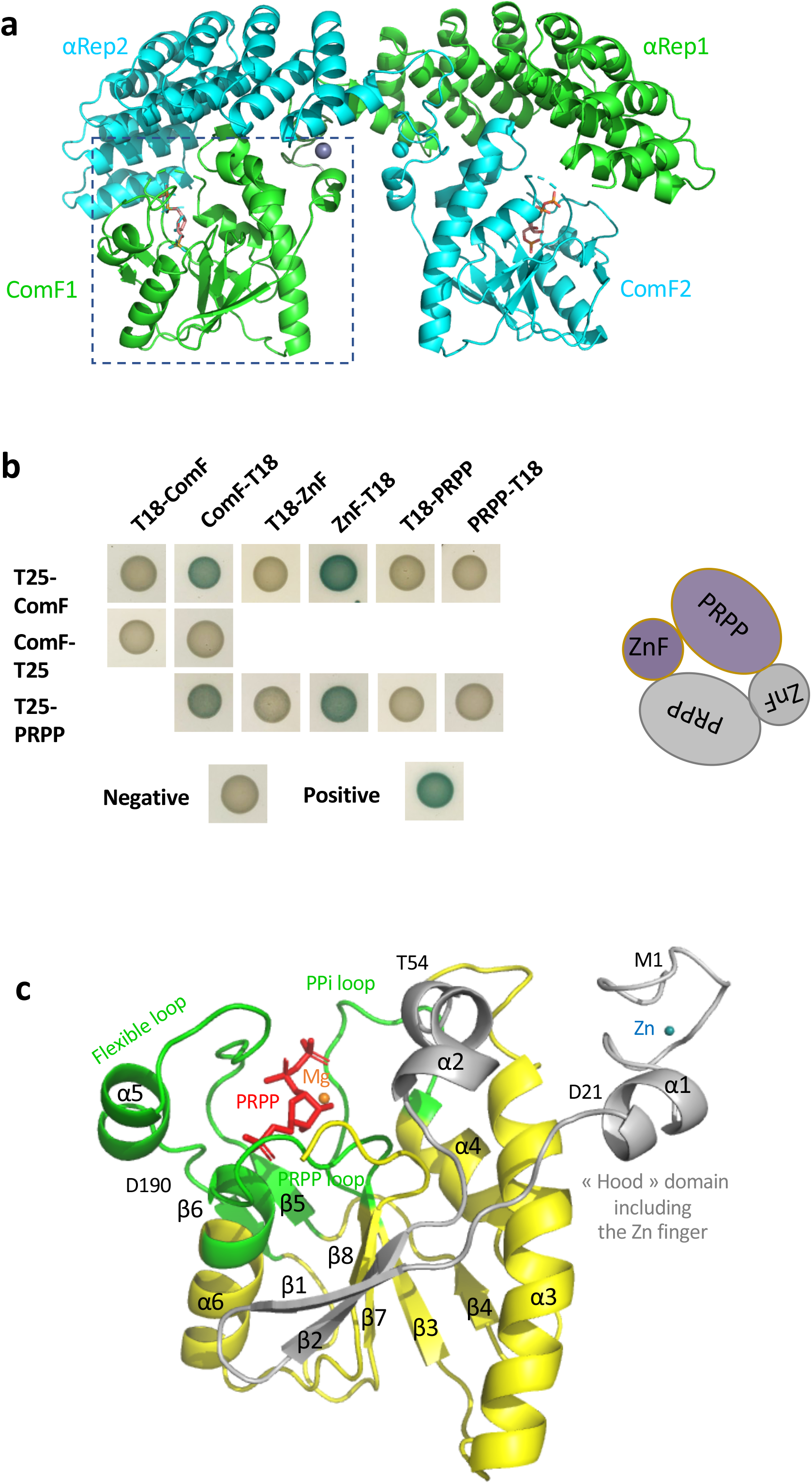
*H. pylori* ComF harbours a PRT and Zn-finger domains. (**a**) Crystal structure of the αRep-HpComF domain-swapped dimer. The two protein fusions are in green and blue. The αRep was evolved against ComF to develop a specific interaction surface. In the crystal packing, each αRep returned to the ComF of another fusion, allowing the crystallization of an artificial dimer. The PRPP co-crystallized with the protein is in sticks, and the Zn^2+^ ion is schematized by a sphere. (**b**) Bacterial two-hybrid assay of *H. pylori* ComF and its N-terminal and C-terminal domains. Representative images of reporter cells grown on plates supplemented with IPTG and X-Gal are shown. (**c**) Structure of ComF. The three loops characteristic of the PRTase fold are in green and the “hood” domain is in grey. The PRPP is in red sticks and the Zn^2+^ and Mg^2+^ ions are schematised by blue and orange spheres, respectively.

The presence of the αRep impeded to conclude from the crystal structure on the possibility of ComF adopting higher order quaternary structures. Most PRT proteins form dimers ^32^. This, together with the dimers described for the *S. pneumoniae* ComFC protein ^18^, prompted us to explore by bacterial two hybrid assays (BacTH) whether the *H. pylori* orthologue could also interact with itself. This was indeed the case (Fig. 5b). To better define the interaction mode, BacTH assays using the separate CTD (residues 26 – 191) and NTD (residues 1 – 25) domains were performed. While no signal above background was obtained when the individual domains were tested for the formation of homodimers, a strong interaction between the PRT and the Zn-finger domains was revealed, suggesting that HpComF could form head to tail dimers (Fig. 5b).

### The zinc-finger domain is required for ComF function

The small NTD of ComF (residues M1– D21), which is part of a larger domain corresponding to the additional and variable “hood” domain of the PRTase family ^32^, is a 4-Cys Zn-finger (in grey in Fig. 5c and Supplementary Figure 4a). The Zn-finger is connected by a five amino acid linker to the rest of the hood domain (residues L22–T54), structured into a small sheet of two β strands followed by a kinked α helix. A Zn^2+^ ion is liganded into the protein via the four cysteine residues (C3, C6, C15 and C18 in pink in Supplementary Figure 4a) which are highly conserved within the ComF(C) family (Supplementary Figure 3).

To explore the role of the zinc-finger, we expressed from either the *rdxA* or the *ureA* loci a ComF version in which cysteines 15 and 18 were replaced by serine (ComF-C15S,C18S), and tested its capacity to complement the transformation phenotype of a *ΔcomF* strain. Unlike the wild-type ComF, the mutant protein could not restore natural transformation (Fig. 1b). Furthermore, mutation of the two cysteines hindered the integration into the bacterial chromosome of a ssDNA delivered by electroporation into the cytosol (Fig. 4a). Zinc-finger domains are most often found in proteins known to bind DNA or RNA ^33^. In particular, 4-Cys zinc-fingers are present in ribosomal proteins or in enzymes involved in DNA replication, recombination and transcription ^34,35^. Unfortunately, attempts to purify ComF versions either mutated in cysteines 15 and 18 or deleted of the zinc-finger domain were unsuccessful, preventing further exploration of its function at the biochemical level. There are, however, several examples of zinc-finger domains that do not participate in nucleic acid binding, but are involved in protein-protein interactions ^36,37^. Interestingly, this is the case for RadA, a DNA helicase implicated in NT of Gram-positive bacteria. While RadA mutated in its 4-Cys domain is still able to bind DNA and to carry out its ATPase and helicase activities, it cannot interact with RecA, thus limiting its D-loop unwinding capacity ^38^.

The closest structural homologue of the ComF zinc-finger is the zinc-finger domain of RecR, a recombination protein (RMSD of 0.8 Å for 22 aligned residues, PDB number 4O6O ^39^ or PDB number 5Z2V ^40^). While the role of the RecR zinc-finger remains to be determined ^41,42^, it has been suggested that it has a structural role in protein folding ^42^. Supporting this possibility, as in our case, the authors were unable to produce soluble forms of *E. coli* or *H. pylori* RecR mutated in its zinc-finger cysteines. While it is tempting to speculate that ComF zinc-finger is involved in the binding of the transforming DNA, we cannot rule out a role of this domain in the interaction of the protein with other NT partners. Further studies are required to define its precise role.

### The PRPP binding domain in necessary for ComF function

The ComF CTD (residues L55– D190, the last E191 is not defined in the electron density) shares the common core of the amidophosphoribosyl transferase type 1 fold (RMSD between 2.6 and 3 Å for 100 to 130 aligned residues, PDB number 5ZGO ^43^ as an example). A central parallel β sheet characteristic of the PRTase core domains is present (β strands ^183^AIA^185, 153^YFLLD^157, 85^LYGIA^89^ and ^113^LKP^115^, in yellow in Fig. 5c), extended by the two β strands of the NTD (^27^KVRVL^31^ and ^34^VSVYS^38^, in grey in Fig. 5c). The three Mg•PRPP-binding loops of the family are present, providing a large hydrogen bonds network with the PRPP (in green in Fig. 5c and Supplementary Figures 3 and 4b). An electron density that can correspond to an Mg^2+^ ion is present close to the PRPP. The “PRPP loop” carries the canonical ^157^DDIITTGTTL^166^ active site signature allowing the binding of the ribose-5-phosphate group of the PRPP (Supplementary Figures 3 and 4b). The most variable “PPi loop” (A^89^ to H^100^) allowing the binding of the PPi group of the PRPP is slightly longer than the standard four amino acids loops. The “flexible loop” (L^120^ to T^144^) closes the pocket of the binding site occupied in our structure by the PRPP (red sticks in Fig. 5c). The presence of all three loops (Supplementary Figures 3) is considered the signature of the PRT family ^32^.

The PRPP-binding domain, present in a large variety of proteins, is known to bind small molecules such as nucleotides or NMPs ^32,44^. We performed differential scanning fluorimetry/thermal shift assays to detect interactions of purified *H. pylori* His_6_-ComF with various potential ligands. Fig. 6a shows that the wild-type protein exhibited a Tm of around 46 ° C. In the presence of AMP the fluorescence maxima observed for the wild-type protein was shifted by +9°C, suggesting the stabilisation of the protein through binding of the nucleotide. Albeit to a lesser extent, ADP addition to ComF also resulted in an increase in melting temperature (Supplementary Table 1). No effect was observed with the triphosphate nucleotide. To confirm that the nucleotide binding was through the PRPP-binding domain, we purified a mutant version of the protein where the threonine165 present in the conserved ^155^LLDDIITTGTTL^166^ motif was replaced by an alanine. ComF T165A had a melting temperature close to that of the wild-type, indicating that the amino acid replacement did not affect significantly the structure of the protein. However, the addition of the nucleotides had a very modest effect on the thermal stability of the protein (Fig. 6b and Supplementary Table 1), confirming the role of the conserved threonine in ligand binding.

**Fig. 6:**
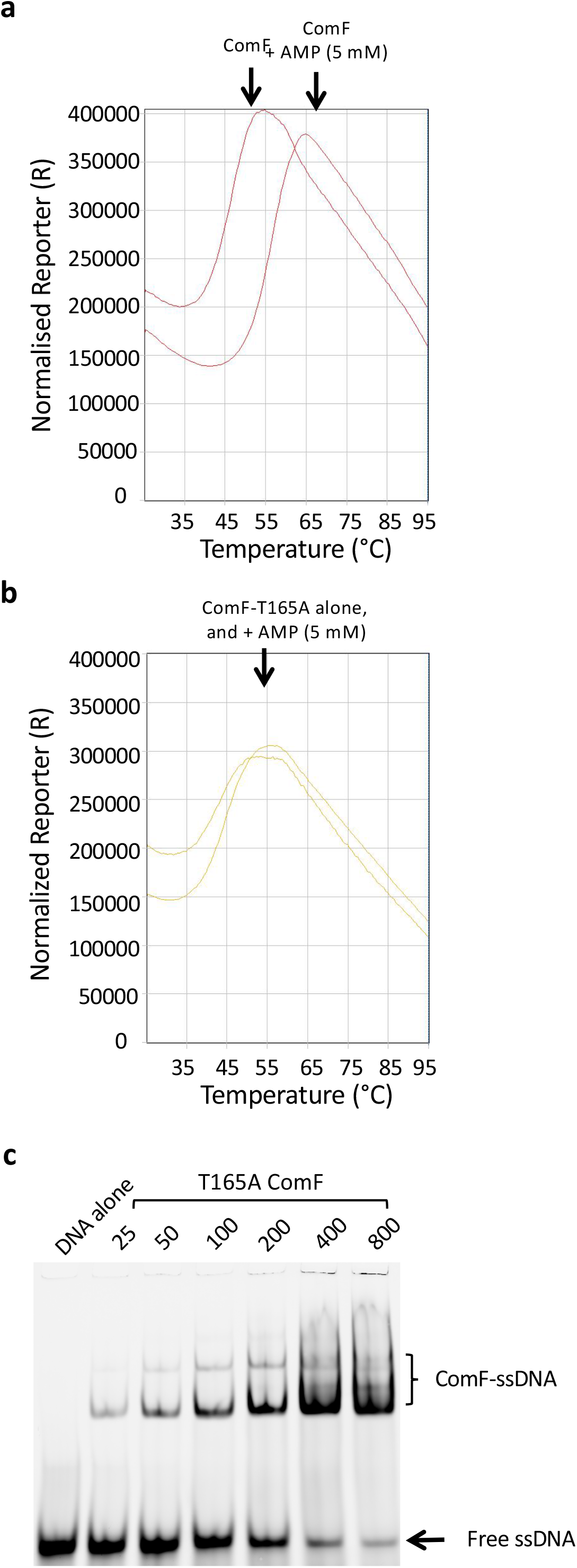
ComF binds to nucleotides through its PRTase motif. Thermal denaturation curves displaying melting temperature of (**a**) ComF and, (**b**) ComF-T165A with or without Adenosine monophosphate. (**c**) Single-stranded DNA binding by ComF-T165A.

To assess the relevance of ComF nucleotide binding capacity in the function of the protein during NT, we expressed in a *ΔcomF* strain either ComF T165A from the *rdxA* locus and its own promoter or ComF-FLAG T165A at the *urea* locus. While ComF T165A restored to a certain extent the transformation capacity, the recombinant frequency was 43-fold lower than that obtained with the wild-type protein expressed in the same conditions (Fig. 1). In the case of the strain expressing ComF-FLAG T165A, the expression of the protein was unable to complement the transformation phenotype of the *ΔcomF* strain (Fig. 1b).

While the presence of a zinc-finger is consistent with an involvement of ComF in the handling of the transforming DNA, the fact that the protein belongs to the PRT family is surprising. ^32^. The majority of PRT proteins are enzymes that catalyse the displacement of pyrophosphate from PRPP by a nitrogen-containing nucleophile ^32^. While there are other PRTases, those belonging to the PRT family are involved in nucleotide synthesis and salvage pathways. Our results (Supplementary Table 1) show that ComF PRT domain is capable of binding not only PRPP but also monophosphorylated nucleotides and, albeit with less affinity, nucleotide di-phosphates. The physiological ligand remains, however, to be determined.

In a few PRT proteins the PRTase capacity to bind PRPP or the nucleotide substrate has been co-opted for regulatory functions as described for two *Bacillus subtilis* regulators of gene expression. PurR binds to DNA operator sequences to repress the expression of purine genes. Binding of PRPP lowers its affinity for the DNA, triggering expression ^45^. PyrR binds regulatory regions of pyrimidine genes transcripts attenuating their expression. Its affinity for the mRNA is regulated by UMP ^46^. We thus asked if the T165A mutation affected ComF affinity for the ssDNA. Even though this substitution abolished the interaction with nucleotides (Supplementary Table 1) it did not significantly affect the binding of ssDNA (Fig. 6c and Supplementary Figure 5).

The experiments presented here do not allow to conclude on whether ComF PRT domain provides a PRTase activity or a regulatory function. An intriguing hypothesis is that the deoxyribomononucleotides released by the degradation of the non-transforming strand ^47^ might regulate ComF capacity to bind DNA. Although ComF T165A is not affected in its DNA affinity (Fig. 6c), it is possible that binding to the wild-type protein of a so far unidentified nucleotide results in a reduced affinity for the transforming DNA. It is worth noting that the T165A mutant, while completely impaired in nucleotide binding (Fig. 6b and Supplementary Table 1), can still partially rescue the transformation phenotype of a *ΔcomF* mutant (Figure 1b), suggesting that nucleotide binding to ComF could provide a fine-tuning mechanism of the transformation process. Such a scenario would be consistent with the recently proposed hypothesis that ComF provides a link between transformation and metabolism ^48^.

## Conclusion

In this study, using *H. pylori* as a model, we sought to unveil the role of ComF, one of the most conserved proteins involved in horizontal gene transfer through NT. Despite the discovery of its essentiality for competence over 30 years ago, the understanding of where and how this protein participates in NT remained elusive. We showed here that ComF is required for at least two different steps in NT. First, ComF facilitates the transport of the tDNA through the cell membrane. Consistent with this finding we found that the protein localises to the inner membrane. Secondly, ComF, which we show has affinity for ssDNA, is involved in the handling of the DNA within the cytosol. We therefore propose that ComF provides a link between these two distinct steps during NT. Our structural studies demonstrated that ComF is composed of two conserved domains, both essential for its *in* vivo activity: a 4-Cys zinc-finger domain and a PRPP-binding domain. While several details of ComF mechanism of action remain to be elucidated, the data presented here shed light on the role of this protein critical for NT in all naturally competent bacteria.

## Methods

### *H. pylori* cultures

*H. pylori* strains are listed in Supplementary Table 2. Cultures were grown under microaerophilic conditions (5% O_2_, 10% CO_2_, using the MAC-MIC system from AES Chemunex) at 37 °C. Blood agar base medium (BAB) supplemented with 10% defibrillated horse blood (AES) was used for plate cultures. Liquid cultures were grown in brain heart infusion media (BHI) supplemented with 10% defibrillated and de-complemented fetal bovine serum (Invitrogen, Carlsbad, CA, USA) with constant shaking (180 rpm). Antibiotic mix containing polymyxin B (0.155 mg/ml), vancomycin (6.25 mg/ml), trimethoprim (3.125 mg/ml), and amphotericin B (1.25 mg/ml) was added to both plate and liquid cultures. Additional antibiotics were added as required: kanamycin (20 μg/ml), apramycin (12.5 μg/ml), and chloramphenicol (8 μg/ml) streptomycin (10 μg/ml) as required.

### Construction of gene variants in *H. pylori*

All oligonucleotides and plasmids used in this work are listed in Supplementary Tables 3 and 4. *H. pylori* 26695 gene sequences were obtained from the annotated complete genome sequence of 26695 deposited at http://genolist.pasteur.fr/PyloriGene/. Gene/locus specific primers (listed in Supplementary Table 3) were used to amplify region of interest by PCR, and fragments were joined together by either classical restriction-ligation method or using sequence- and ligation-independent cloning (SLIC). Different protein tags and mutations in the genes were introduced using SLIC or site directed mutagenesis, respectively. All the plasmids generated were verified by DNA sequencing (listed in Supplementary Table 4). The knock-out /Knock-in cassette were introduced into *H. pylori* by natural transformation. Their correct integration in *H. pylori* genome was confirmed by PCR using locus and gene specific primers. Verified strains (Table S2) were stored at -80 °C in BHI media supplemented with 12.5 % glycerol. The details of different constructions generated in this study are given below.

### Construction of *hp1473* null mutants *in H. pylori*

To generate *hp1473* locus disrupted by a non-polar chloramphenicol cassette (*hp1473::Cm*), *hp1473* locus was amplified using gene specific primers (hp1473F and hp1473R) and ligated to blunt pjET1.2 vector to generate pJet1.2-*hp1473*. PCR fragments generated by amplification of this plasmid (using primers 1473 inverse F and 1473 inverse R), and non-polar chloramphenicol (using primers KpnI-Cm-for and BamHI-Cm-rev) resistance cassette were digested using KpnI and BamHI and then ligated to generate the knockout cassette p978 (pJet1.2-*hp1473::Cm*).

To generate *hp1473* locus disrupted by a non-polar apramycin cassette (*hp1473::apramycin*), *hp1473* locus was amplified using primers Op853 and Op854 and ligated to pjET1.2 vector amplified using Op855-Op856 to generate pJet1.2-*hp1473*. The PCR fragments generated by amplification of pJet1.2-*hp1473* (using primers Op859 and Op860) and a non-polar apramycin resistance cassette (using primers Op857 and Op858) were ligated using SLIC to generate p1699.

### Ectopic expression of *hp1473* variants

*hp1473* locus (+ 152 bp upstream sequence) was amplified using Op5 and Op6 and inserted in *rdXA::km* cassette present in the plasmid p1175 (amplified using Op3 and Op4) using SLIC to generate plasmid p1176. T165A and C15SC18S mutations in the hp1473 coding region were introduced by PCR using mutagenic primers (op13+ Op14 and Op302 + Op303 respectively). The native *hp1473* locus was disrupted using p978. For expression using the *urea* promoter *hp1473* loci, wild-type or containing T165A or C15SC18S point mutations were amplified using Op611 and Op612 (containing the sequence for Flag tag) was ligated with p1088 (amplified using Op613 and Op614) containing the Promoter-UreA-Cm cassette using SLIC to generate p1672, p1674 and p1676 respectively. These plasmids were used to transform *H. pylori* followed by disruption of native *hp1473* locus was using p1699.

### Determination of transformation frequencies

Natural transformation frequencies were determined as described ^49^. Briefly, total chromosomal DNA (200 ng) from a streptomycin resistant but otherwise isogenic strain was incubated overnight with exponentially growing *H. pylori* cells (optical density of 4.0 at 600 nm), on solid medium. Next day, serial dilutions of *H. pylori* were spread on plates with and without streptomycin (10 μg/ml). Transformation frequencies after electroporation were determined as described ^10^. Briefly, electro-competent cells were prepared by treating *H. pylori* cells (optical density of 10 OD/ml at 600 nm) with ice-cold Glycerol 15 % + Sucrose 9 %. 50 µl of electro-competent cells mixed with 1 µg of 139-mer or 75-mer-ssDNA (Supplementary Table 3) carrying A128G mutation in the *hp1197* gene were electroporated at 2.5 kV cm^-1^ and 25 µF. The cells were mixed with 100 µl BHI, and 50 µl cells were spotted on BAB plates. Next day, serial dilutions of *H. pylori* cells were plated on plates with or without streptomycin (10 μg/ml). The transformation frequencies were calculated as the number of streptomycin resistance colonies per recipient colony-forming unit. *P* values were calculated using the Mann–Whitney *U* test on GraphPad Prism software.

### Fluorescence microscopy experiments

Microscopy experiments were performed as described earlier ^10,12^. Fluorescent dsDNA (408 bp) was prepared by amplification of *hp1197* locus from 26695 gDNA (100 ng) using primers 1197-5’ and 1197-3’ (0.5 μM each), 250 μM of dNTP mix, 5 Units of ExTaq enzyme (Takara) supplemented with 10 μM of ATTO-550-aminoallyl-dUTP (Jena bioscience). PCR elongation was performed at 72 °C (2 min per kb) and the amplified products were purified by illustra GFX purification kit (GE Healthcare Little Chalfont, UK).

Exponentially growing *H. pylori* cells were incubated with fluorescent DNA (200 ng) for 7 min at 37 °C, the unbound DNA was washed the bacteria were re-suspended in BHI, covered with low melting agarose (1.4 %) supplemented with 10 % fetal bovine serum and were observed under live conditions [gas mixture (10 % CO2, 3 % O2), humidity (90 %)] at 37 °C for 3 h. Alternatively, the bacteria were fixed with 4 % formaldehyde (90 mins at 4 °C) followed by quenching with 100 mM Glycine. All the images were captured with 60X objective using inverted Nikon A1R confocal laser scanning microscope system. The images were processed and analyzed using NIS-element software (Nikon Corp., Tokyo, Japan) and ImageJ software. The percentage of bacteria with DNA foci was calculated as the number of bacteria with DNA foci over total number of bacteria counted in at least two independent biological replicates. To monitor the time dependent stability DNA foci, the total number of bacteria with DNA foci at t=15 mins were considered as 100%. The volumes of internalised DNA in GFP expressing bacteria were estimated by 3-D image analysis performed using Volocity software (Perkin Elmer, Waltham, USA).

### Subcellular fractionation

Subcellular fractions of exponentially growing *H. pylori* were collected by differential centrifugation and detergent mediated solubilisation as described earlier ^10^. Briefly, 100 ml cell pellet was re-suspended in buffer A (10 mM Tris-HCl, pH 7.5, 1mM DTT, 1X protease inhibitor cocktail) followed by lysis by sonication. Total extracts were centrifuged 14,000 rpm for 15 minutes. The supernatant containing the soluble fraction was collected after ultracentrifugation at 45,000 rpm for 45 min of the total extract. The pellet containing the membrane fractions was re-suspended in buffer B (10 mM Tris-HCl, pH 7.5, 1mM DTT, 1X protease inhibitor cocktail, 1% N-Lauroylsarcosine). The supernatant containing the inner membrane fractions were collected after ultracentrifugation at 45,000 rpm for 45 min. The presence of the proteins in the various fractions was monitored by immunoblotting.

### Western blots

The fractionation samples were resolved on a 15% SDS-PAGE, transferred on a nitrocellulose membrane. The membrane was blocked with 2% BSA prepared in PBST (1X PBS + 0.03% Tween 20). Blots were probed with either a mouse monoclonal anti-Flag antibody (1: 5000 dilution, Sigma Aldrich), rabbit anti-MotB antibody (1: 2500 dilution) (kind gift from Dr. Ivo Boneca, Pasteur Institute) or rabbit anti-HpComF antibody from our laboratory collection. The blots were then probed with Advansta fluorescently labeled secondary antibodies IR700 and IR800 respectively. The imaging was done using Odissey Clx imaging system.

### *E. coli* cultures

*Escherichia coli* strains used for cloning, protein overexpression and purification were cultured in Luria–Bertani (LB) broth, or LB agar plates supplemented with the required antibiotics [ampicillin (100 μg/ml), kanamycin (50 μg/ml), apramycin (50 μg/ml), or chloramphenicol (34 μg/ml)].

### Protein samples preparation

Cloning of *comF* (*Hp1473*) coding region was performed using genomic DNA from *H. pylori* strain 26695 as template for PCR. Six histidine codons were added at the 5’ end during the PCR process. The fragment was inserted into the *Nde*I-*Xho*I sites of pET21a vector (Novagen). Site directed mutagenesis was performed using the resulting pET21:ComF-6His plasmid as a template and non-overlapping oligonucleotides phosphorylated in 5’ (Eurofins), to construct the *HpComF*^T165A^ mutant. The fusion of *comF* with an αRep protein (named B2), for its structural study, is described in ^31^.

Expression of ComF or its mutant forms in BL21(DE3) Gold strain was performed in 800 ml 2xYT o/n at 37°C after induction with 0.5 mM IPTG (Sigma). Cells were harvested by centrifugation, resuspended in buffer 500 mM NaCl, 20 mM Tris-HCl (pH 7.5), 5% glycerol for ComF-6His constructs, or in buffer 1 M NaCl, 100 mM Tris-HCl (pH 8), 100 µM TCEP for B2-ComF-6His (SeMet labelled according to the protocol described in ^50^ and stored at –20°C. Cell lysis was completed by sonication (probe-tip sonicator Branson). After centrifugation at 20,000 g and 8°C for 30 min, the proteins were purified on a Ni-NTA column (Qiagen Inc.), eluted with imidazole. ComF-6His and B2-ComF-6His were then desalted up to 100 mM NaCl and loaded respectively onto a Heparin and a MonoQ column (Amersham Pharmacia Biotech) and eluted with a gradient of NaCl (from 100 mM to 1 M). The proteins were desalted up to 200 mM NaCl and concentrated using Vivaspin 5,000 or 30,000 nominal molecular weight limit cut-off centrifugal concentrators (Sartorius), respectively, aliquoted, flash frozen in liquid nitrogen and stored at -80°C, or dialyzed in a 50% (vol/vol) glycerol buffer for storage at -20°C.

### Electrophoretic mobility shift assay

DNA biding assays were performed by incubating indicated concentrations of proteins with fixed concentrations of Cy5 labelled DNA (Supplementary Table 3) in binding buffer (10 mM Tris-HCl pH 7.5, 50 mM KCl, 1 mM DTT, 0.1 µg/µl BSA) in cold room for 30 min. The nucleoprotein complexes were separated using native TBE-PAGE (6%). The gels were visualized by using Typhoon. The depletion in substrate DNA was quantified using ImageJ by considering DNA without protein as 100%.

### Crystal structure determination

SeMet modified B2-HpComF-6His (12.5 mg/ml) was incubated with PRPP (3 mM) and MgCl_2_ (5 mM) at 4°C. Crystals were grown in hanging drops by mixing the protein with reservoir solution in a 1:1 ratio. Crystals appeared after 5 days at 4°C in 0.2 M Tri-potassium citrate + 18% PEG 3350. Glycerol cryo-protected crystals (two steps at 15 and 30%) were flash frozen in liquid nitrogen.

Diffraction data and refinement statistics are given in Table 1. Crystallographic data were collected at the selenium peak wavelength on the PROXIMA-2A from Synchrotron SOLEIL (Saint-Aubin, France) and processed with XDS ^51^ through XDSME (https://githubcom/legrandp/xdsme). Diffraction anisotropy was corrected using the STARANISO server (http://staraniso.globalphasing.org). The structure was solved by the SAD phasing method at 2.5 Å resolution using SHELX C/D ^52^ to locate the 12 heavy atom sites, PHASER ^53^ to determine the initial phases and PARROT ^54^ to improve the phases by density modification, through the CCP4 program suite ^55^. The construction of the model was initiated using Buccaneer ^56^ and refined with the BUSTER using TLS and NCS restraints ^57^. The model was corrected and completed using COOT ^58^. The presence of a Zn^2+^ ion in the B2-HpComF structure was demonstrated by an energy scan performed on the crystals at the beamline (energy peak at 9.664 keV). Exploration of the 3D structures was performed using the following tools: Dali server ^59^, I-TASSER ^60^ and SWISS-MODEL servers ^61^ and PyMOL Molecular Graphics System (http://www.pymol.org).

### Bacterial Two-Hybrid assays

The Bacterial Two-Hybrid test was used to probe the interactions between proteins ^62^. The full-length ComF encoding sequence was fused to T18, at the C-terminal and N-terminal ends, T18-ComF (pUT18C vector) and ComF-T18 (pUT18 vector), respectively. The same strategy has been used for ZnF and PRPP, the N-terminal and C-terminal domains of ComF, respectively. Plasmids encoding T25-ComF, ComF-T25 and T25-PRPP were constructed using the pKT25 and pKNT25 vectors.

Plasmids encoding T18 and T25 fusion proteins were co-transformed in *E. coli* strain BTH101 and transformants were selected in Luria-Bertani agar plates containing kanamycin and ampicillin at 30°C. Colonies were then spotted on plates containing kanamycin, ampicillin, IPTG and X-gal, incubated at 30°C and stored at RT to follow the appearance and evolution of the blue colour.

### Differential Scanning Fluorimetry/Thermal shift assay

Purified protein (10.5 µg) was incubated with different analytes in reaction buffer (20 mM Tris-Cl, pH 7.5, 200 mM NaCl, 5X Sypro Orange). The temperature of the reaction mixture was raised from 25 ° C to 95 ° C. Shift in the fluorescence due to binding of the Sypro-Orange dye as the hydrophobic patches of the protein were exposed due to denaturation of the protein was recorded. The fluorescence maxima observed was used to calculate the approximate melting temperature of the protein in native conditions and in presence of the analyte.

## Supporting information

Supplementary movie 1 - Wild-type

## Data availability

The atomic coordinates and structure factors of B2-HpComF have been deposited at the Brookhaven Protein Data Bank under the accession number 7P0H. All the other data are available in the main text or the supplementary materials.

## Acknowledgements

We thank the beamline staff for assistance and advice during data collections at Synchrotron SOLEIL (Saint-Aubin, France; beamline Proxima 2). We thank Christopher Corbinais and Mariano Prado-Acosta for the construction of the initial *comF* mutants. This work has benefited from the I2BC Macromolecular interactions measurements and Crystallization Platforms. Financial support for this work was provided by the Indo-French Centre for Promotion of Advanced research (CEFIPRA) grant 5203-5 (JPR, PPD), Agence Nationale de la Recherche grant ANR-19-CE12-0003-01 (JPR), French Infrastructure for Integrated Structural Biology (FRISBI) grant ANR-10-INSB-05-01 (SQC, RG), Région Ile de France grant DIM1Health (JPR, PPD), Commissariat à l’Energie Atomique (JPR, AMDG, XV, JD), Centre National de la Recherche Scientifique (SQC, SM), *Enhanced Eurotalents* fellowship programme (CEA/EU) (PPD) and Collectivité Régionale de Martinique (LC)

## Author contributions

Conceptualisation: SQC, JPR, PPD

Methodology: SQC, SM, PPD, AMDG, XV, JPR

Formal analysis: HW, JV, RG, SQC, JPR

Investigation: PPD, LC, SK, AMDG, SM, JD, XV, SQC, PL

Data curation: LC, HW, PL

Writing - Original Draft: PPD, SQC, JPR

Writing – Review and Editing: PPD, SQC, JPR. All authors read and approved the manuscript.

Visualisation: PPD, SK, AMDG, SM, HW, SQC, JPR

Supervision: JPR, SQC

Project administration: JPR

Funding acquisition: JPR, SQC

## Competing interests

The authors declare that they have no competing interests.

## Notes

### Competing Interest Statement

The authors have declared no competing interest.

