## Supplementary material for "ComF is a key mediator in single-stranded DNA transport and handling during natural transformation"

**Other Supplementary Information for this manuscript includes the following:**

Movies S1 to S3

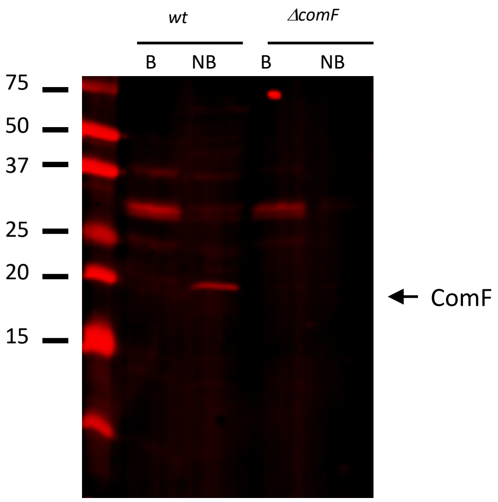


**Supplementary Figure 1.**

**ComF detection by immunoblotting.** Total extracts from wild-type (*wt*) and *comF* mutant (*ΔcomF*) *H. pylori* strains. Samples were either boiled (B) or not boiled (NB) prior gel loading. ComF is specifically detected in the NB condition.

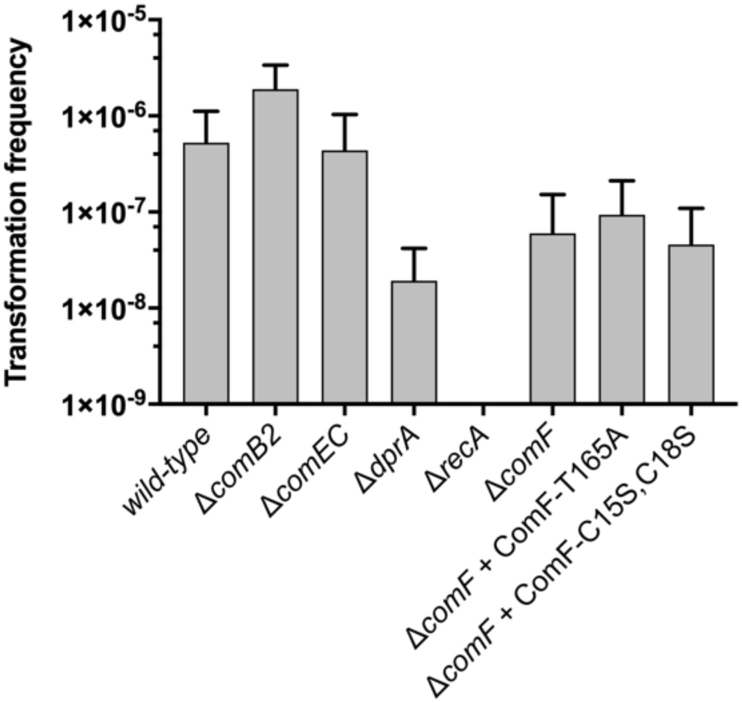


**Supplementary Figure 2**

**Transformation frequencies after electroporation** with a chemically synthesized single-stranded DNA (139 -mer) coding for streptomycin resistance as donor DNA. Bars correspond to the average and standard deviation from at least two independent biological experiments.

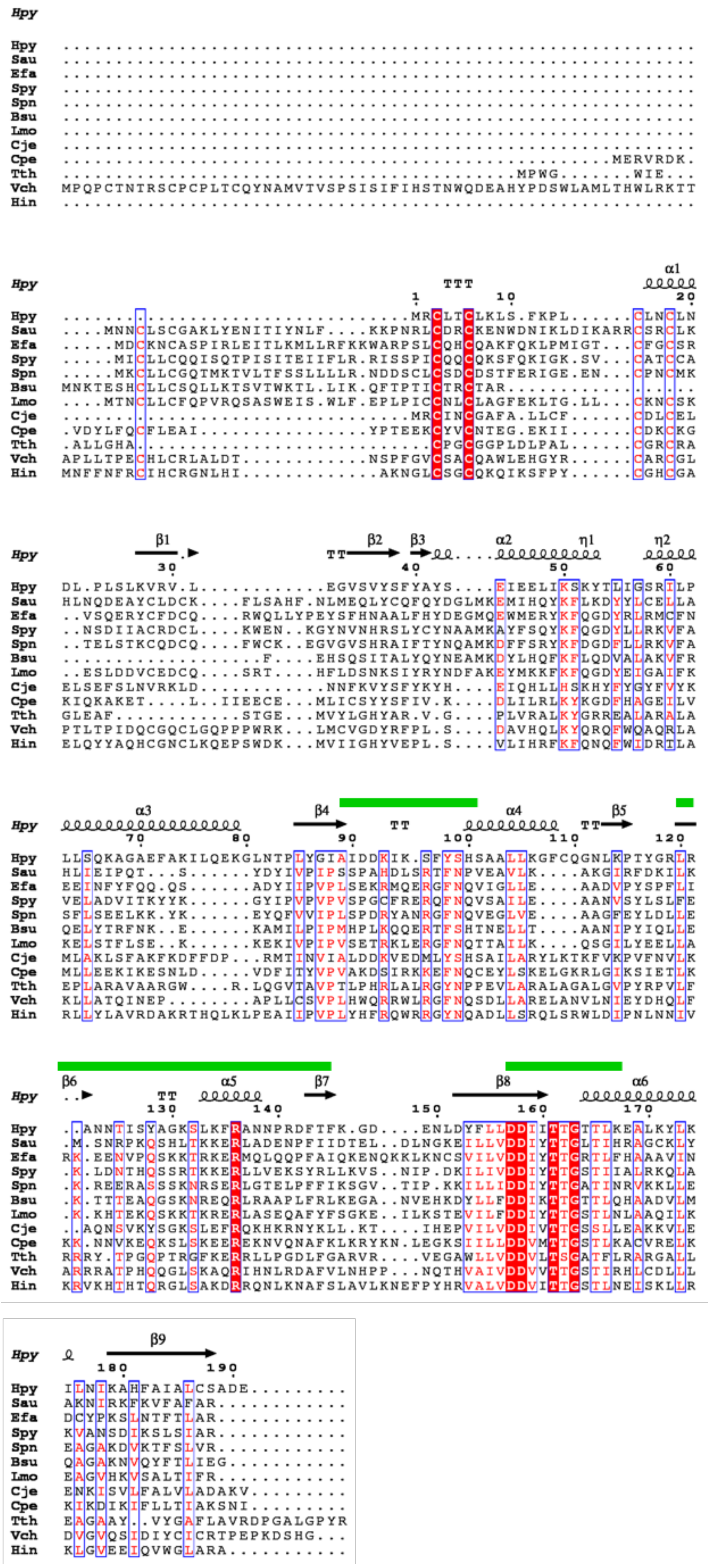
Hpy
Hpy ..........................................................
Sau ..........................................................
Efa ..........................................................
Spy ..........................................................
Spn ..........................................................
Bsu ..........................................................
Lmo ..........................................................
Cje ..........................................................
Cpe .....................................................MERVRDK.
Tth .........................................MPWG.......WIE...
Vch MPQPCTNTRSCPCPLTCQYNAMVTVSPSISIFIHSTNWQDEAHYPDSWLAMLTHWLRKTT
Hin ..........................................................
α1
Hpy TTT
1 10 20
Hpy ...............................MRCLTCLKLS.FKPL......CLNCLN
Sau ....MNNCLSCGAKLYENITIYNLF....KKPNRLCDRCKENWDNIKLDIKARRCSRCLK
Efa .....MDCKNCASPIRLEITLKMLLRFKKWARPSLCQHCQAKFQKLPMIGT...CFGCSR
Spy .....MICLLCQQISQTPISITEIIFLR.RISSPICQQCQKSFQKIGK.SV...CATCCA
Spn .....MKCLLCGQTMKTVLTFSSLLLLR.NDDSCLCSDCDSTFERIGE.EN...CPNCMK
Bsu MNKTESHCLLCSQLLKTSVTWKTL.LIK.QFTPTICTRCTAR................
Lmo ....MTNCLLCFQPVRQSASWEIS.WLF.EPLPICCNLCLAGFEKLTG.LL...CKNCSK
Cje ...............................MRCINCGAFA.LLCF.....CDLCEL
Cpe .VDYLFQCFLEAI...............YPTEEKCYVCNTEG.EKII......CDKCKG
Tth .ALLGHA.....................CPGCGGPLDLPAL.....CGRCRA
Vch APLLTPECHLCRLALDT...........NSPFGVCSACQAWLEHGYR......CARCGL
Hin MNFFNFRCIHCRGNLHI.............AKNGLCSGCQKQIKSFPY......CGHCGA
β1 β2 β3 α2 η1 η2
Hpy TT
30 40 50 60
Hpy DL.PLSLKVRV.L...........EGVSVYSFYAYS....EIEELIKSKYTLIGSRILP
Sau HLNQDEAYCLDCK....FLSAHF.NLMEQLYCQFQYDGLMKEMIHQYKFLKDYYLCELLA
Efa ..VSQERYCFDCQ....RWQLLYPEYSFHNAALFHYDEGMQEWMERYKFQGDYRLRMCFN
Spy ..NSDIIACRDCL....KWEN..KGYNVNHRSLYCYNAAMKAYFSQYKFQGDYLLRKVFA
Spn ..TELSTKCQDCQ....FWCK..EGVGVSHRAIFTYNQAMKDFFSRYKFDGDFLLRKVFA
Bsu ...................F....EHSQSITALYQYNEAMKDYLHQFKFLQDVALAKVFR
Lmo ..ESLDDVCEDCQ....SRT....HFLDSNKSIYRYNDFAKEYMKKFKFQGDYEIGAIFK
Cje ELSEFSLNVRKLD...........NNFKVYSFYKYH....EIQHLLHSKHYFYGYFVYK
Cpe KIQKAKET....L..IIEECE...MLICSYYSFIV.K....DLILRLKYKGDFHAGEILV
Tth GLEAF...........STGE...MVYLGHYAR.V.G....PLVRALKYGRREALARALA
Vch PTLTPIDQCGQCLGQPPPWRK...LMCVGDYRFPL.S....DAVHQLKYQRQFWQAQRLA
Hin ELQYYAQHCGNCLKQEPSWDK...MVIIGHYVEPL.S....VLIHRFKFQNQFWIDRTLA
α3 β4 α4 β5
Hpy TT TT
70 80 90 100 110 120
Hpy LLSQKAGAEFAKILQEKGLNTPLYGIAIDDKIK.SFYSHSAALLKGFCQGNLKPTYGRLR
Sau HLIEIPQT...S......YDYIVPIPSSPAHDLSRTFNPVEAVLK....AKGIRFDKILK
Efa EEINFYFQQ.QS......ADYIIPVPLSEKRMQERGFNQVIGLLE....AADVPYSPFLI
Spy VELADVITKYYK......GYIPVPVPVSPGCFRERQFNQVSAILE....AANVSYLSLFE
Spn SFLSEELKK.YK......EYQFVVIPLSPDRYANRGFNQVEGLVE....AAGFEYLDLLE
Bsu QELYTRFNK..E......KAMILPIPMHPLKQQERTFSHTNELLT....AANIPYIQLLE
Lmo KELSTFLSE..K......KEKIVPIPVSETRKLERGFNQTTAILK....QSGILYEELLA
Cje MLAKLSFAKFKDFFDP...RMTINVIALDDKVEDMLYSHSAILARYLKTKFVKPVFNVLK
Cpe MLLEEKIKESNLD.....VDFITYVPVAKDSIRKKEFNQCEYLSKELGKRLGIKSIETLK
Tth EPLARAVAARGW....R.LQGVTAVPTLPHRLALRGYNPPEVLARALAGALGVPYRPVLF
Vch KLLATQINEP........APLLCSVPLHWQRRWLRGFNQSDLLARELANVLNIEYDHQLF
Hin RLLYLAVRDAKRTHQLKLPEAIIPVPLYHFRQWRRGYNQADLLSRQLSRWLDIPNLNNIV
β6 α5 β7 β8 α6
Hpy ..
TT
130 140 150 160 170
Hpy ..ANNTISYAGKSLKFRANNPRDFTFK.GD....ENLDYFLLDDIITTGTTLKEALKYLK
Sau .M.SNRPKQSHLTKKERLADENPFIIDTEL..DLNGKEILLVDDIYTTGLTIHRAGCKLY
Efa RK.EENVPQSKKTRKERMQLQQPFAIQKENQKKLKNCSVILVDDIYTTGRTLFHAAAVIN
Spy .K.LDNTHQSSRTKKERLLVEKSYRLLKVS..NIP.DKILIVDDIYTTGSTIIALRKQLA
Spn .K.REERASSSKNRSERLGTELPFFIKSGV..TIP.KKILLIDDIYTTGATINRVKKLLE
Bsu .K.TTTEAQGSKNREQRLRAAPLFRLKEGA..NVEHKDYLLFDDIKTTGTTLQHAADVLM
Lmo .K.KHTEKQSKKTKRERLASEQAFYFSGKE..ILKSTEVILFDDIYTTGSTLNLAAQILK
Cje ..AQNSVKYSGKSLEFRQKHKRNYKLL.KT....IHEPVILVDDIVTTGSSLLEAKKVLE
Cpe KK.NNVKEQKSLSKEERENVQNAFKLKRYKN.LEGKSIILLDDVMTTGSTLKACVRELK
Tth RRRY.TPGQPTRGFKERRLLPGDLFGARVR....VEGAWLLVDDVLTSGATFLRARGALL
Vch ARRRATPHQQGLSKAQRIHNLRDAFVLNHPP...NQTHVAIVDDVVTTGSTIRHLCDLLL
Hin KRVKHTHTQRGLSAKDRRQNLKNAFSLAVLKNEFPYHRVALVDDVITTGSTLNEISKLLR
β9
Hpy
180 190
Hpy ILNIKAHFAIALCSADE.........
Sau AKNIRKFKVFAFAR............
Efa DCYPKSLNTFTLAR............
Spy KVANSDIKSLSIAR............
Spn EAGAKDVKTFSLVR............
Bsu QAGAKNVQYFTLIEG...........
Lmo EAGVHKVSALTIFR............
Cje ENKISVLFALVLADAKV.........
Cpe KIKDIKIFLLTIAKSNI.........
Tth EAGAAY..VYGAFLAVRDPGALGPYR
Vch DVGVQSIDIYCICRTPEPKDSHG...
Hin KLGVEEIQVWGLARA...........

**Supplementary Figure 3.**

**Sequence alignment of the ComFC proteins from gram(-) and gram(+) bacteria**

The multialignment was generated by Clustalw2 (32). The figure was generated using ESPRIPT (33). The secondary structure elements of ComF are in black on the top of the multialignment. Grey and green bars respectively localize the Hood domain and the three PRTase-loops. The 4 cysteine residues of the Zn-finger are indicated by orange stars. Hpy: *Helicobacter pylori*, Sau: *Streptococcus aureus*, Efa: *Enterococcus faecalis*, Spy: *Streptococcus pyrogenis*, Spn: *Streptococcus pneumonia*, Bsu: *Bacillus subtilis*, Lmo: *Listeria monocytogenes*, Cje: *Campylobacter jejuni*, Cpe: *Clostridium perfringens*, Tth: *Thermus thermophilus*, Vch: *Vibrio cholerae*, Hin: *Haemophilus influenzae*.

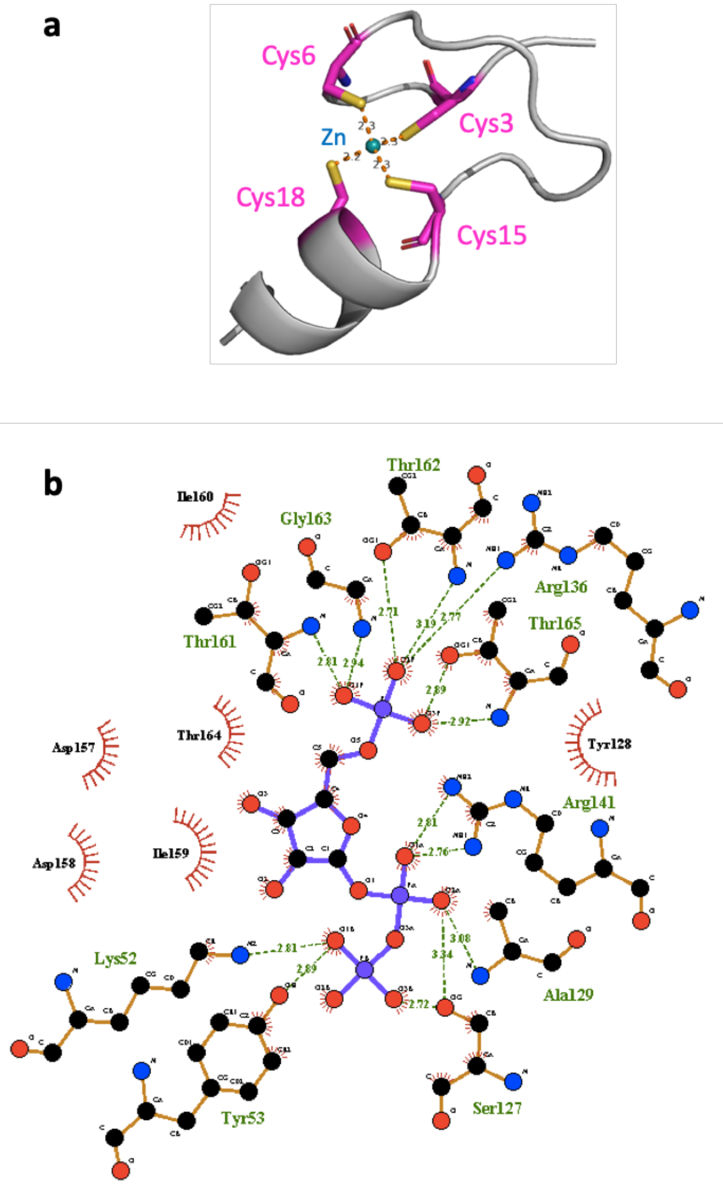


**Supplementary Figure 4.**

**Zoom on the ligand interfaces of HpComF for $Zn^{+2}$ and PRPP**. (**A**) the Zn-finger of ComFC is composed of 4 cysteines (**B**) ComF binds the PRPP (in blue sticks) through hydrogen bonds (names of involved residues in green) and hydrophobic forces (names of involved residues in black). The two PRPP and flexible loops are involved. The 9 residues of the PRPP loop are the characteristic motif of the PRTase family. 5 residues of the flexible loop can close the active site pocket to sequester the PRPP. Two other residues (K52 and Y53) located between helixes α2 and α3 are also involved in the binding of the PRPP.

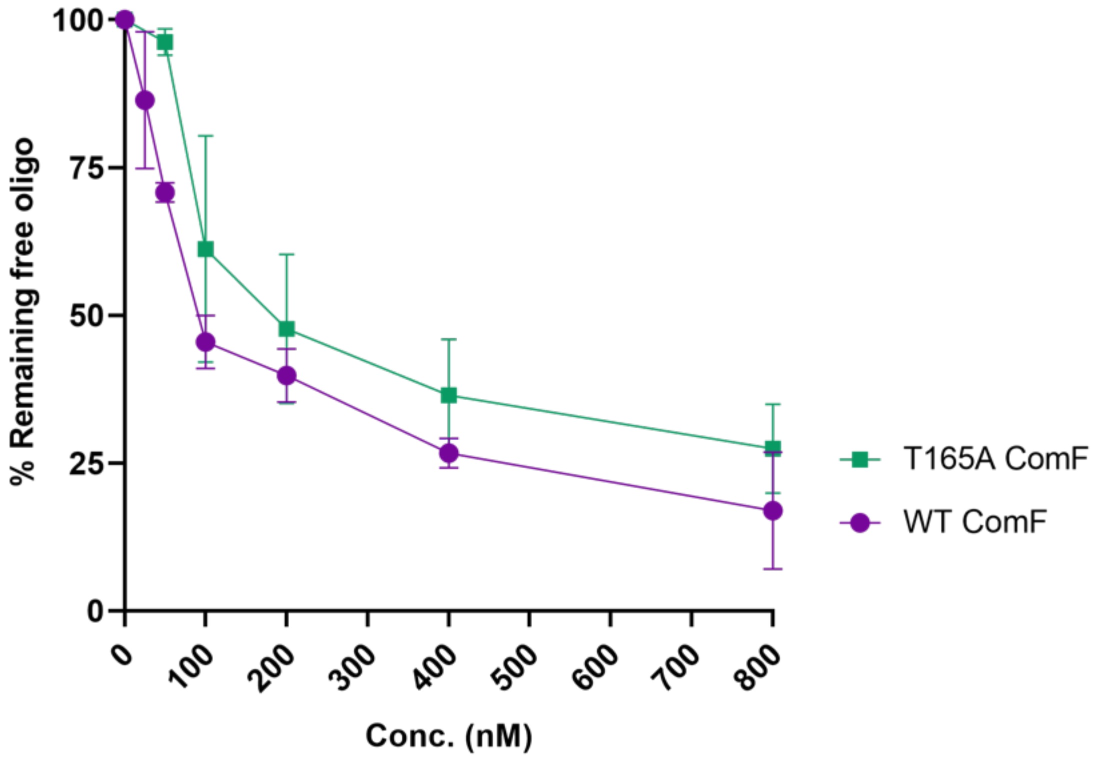


**Supplementary Figure 5.**

**Comparison of DNA binding affinities of WT ComF with ComF T165A.** No significant difference was observed between the binding of WT and T165A ComF proteins to ssDNA. Quantification of three independent electrophoretic mobility shift assays was performed using Image Studio software.

**Table S1.**

**Table S1.**

Approximate melting temperature ($^{o}$C) of wild-type ComF and ComF-T165A with or without added ligand

<

| Ligand | Wild-type ComF | ComF-T165A |
|---|---|---|
| AMP (0) | 45.67 | 42.14 |
| AMP (5mM) | 55.13 | 43.67 |
| AMP (10 mM) | 55 | 45.2 |
| ADP (0) | 45 | 41.21 |
| ADP (5mM) | 51.43 | 44.56 |
| ADP (10 mM) | 51.59 | 42.9 |
| ATP (0) | 46.7 | 41.36 |
| ATP (5mM) | 45 | 41.21 |
| ATP (10 mM) | 45.76 | 41.48 |
| Ribose-5-Phosphate (0) | 44.85 | 41.33 |
| Ribose-5-Phosphate (5 mM) | 48.24 | 41.98 |
| Ribose-5-Phosphate (10 mM) | 48.24 | 41.83 |

**Table S2.**

**Strains used in this study**

| **Strain** | **Genotype** | **Source** |
|---|---|---|
| LR1 | 26695 | |
| LR133 | 26695 *strep*$^{R}$ | Lab. collection |
| LR293, LR294 | 26695 *recA::Cm* | Lab. collection |
| LR827, LR828 | 26695 *dprA::Cm* | Lab. collection |
| LR768, LR769 | 26695 *comB2::Cm* | Lab. collection |
| LR776, LR777 | 26695 *comEC::Km* | Lab. collection |
| LR887 | 26695 *pUreA-GFPmut2-Km* | Lab. collection |
| LR901, LR902 | 26695 *pUreA-GFPmut2-Km comEC::Cm* | Lab. collection |
| LR982 | 26695 *pUreA-GFPmut2-Km hp1473::Cm* | This work |
| LR762 | 26695 *hp1473::Cm* | This work |
| LR965 | 26695 *hp1473::Cm rdxA ::hp1473-Km* | This work |
| LR1038 | 26695 *hp1473::Cm rdxA ::hp1473-FLAG-Km* | This work |
| LR1000 | 26695 *hp1473::Cm rdxA ::hp1473-T165A-Km* | This work |
| LR1051 | 26695 *hp1473::Cm rdxA ::hp1473-C15SC18S-FLAG-Km* | This work |
| LR1209 | 26695 *pUreA-hp1473 flag-Cm hp1473::Apra* | This work |
| LR1211 | 26695 *pUreA-hp1473 flag-Cm hp1473::Apra* | This work |
| LR1213 | 26695 *pUreA-hp1473 flag-Cm hp1473::Apra* | This work |

### Table S3.
### Oligonucleotides used in this study

| Name | Sequence (5'-3') | Description |
|---|---|---|
| 1473 F | ATGCGCTGTTTAACCTGTTTG | hp1473 forward |
| 1473 R | TCATTCATCCGCGCTGCAAAG | hp1473 reverse |
| 1473 inverse R | CGGGGTACC GATTTTCACAAACTCTGCACC | -- |
| 1473 inverse F | CGCGGATCC GGCGTTTAAGGGCTAATAATGC | -- |
| Op3 | GTAATTTTTCTATGCCTTGGTTTTCTTATTCCTCCTAGTTAGTCA GGTACC | HP1473 954 |
| Op4 | CTTTGCAGCGCGGATGAATGAATGGCTAAAATGAGAATATCAC C | HP1473 KanR |
| Op13 | GCCCTAAAAGAAGCCCTAAAATACCTTAAAAC | TA fw |
| Op14 | GTTTTAAGGTATTTTAGGGCTTCTTTTAGGGCGGTGCCGGTGG TG | TA rev |
| Op302 | GCTTTCT TTT AAG CCT CTT TCC CCA AAT TCC TTGAACGATTTGCCCTTAAGCTTAAAGG | C15SC18S F |
| Op303 | CCTTTAAGCTTAAGGGCAAATCGTTCAAGGAATTTGGGGAAAG AGGCTTAAAAGAAAGC | C15SC18S R |
| Op245 | AATCACCTGCATTAGTGGAATGCCCTCAAAGGAGAGGGGTTT GTACTAGGGTTTATACGACTACCCCTAGAAAGCCTAACTCGGC TTTAAGAAAGGTTGCCAAAGTTCGTTTGACCAGTAAATTTGAA GTGATCAGTTA | Streptomycin resistant (139-mer) ssDNA for electroporation |
| Op247 | GAGGGGTTTGTACTAGGGTTTATACGACTACCCCTAGAAAGCC TAACTCGGCTTTAAGAAAGGTTGCCAAAGTTC | Streptomycin resistant (75-mer) ssDNA for electroporation |
| Op611 | CTTTAAGAATAGGAGAATAAGGAATTC**ATG**CGCTGTTTAACCT GTTTGAAGC | For PureA-ATG/ComFC |
| Op612 | TTACTTATCGTCGTCATCCTTGTAATCTTCATCCGCGCTGCAAA GCGCG | Rev ComFC-FLAG*taa |
| Op613 | GCTTCAAACAGGTTAAACAGCGCATGAATTCCTTATTCTCCTAT TCTTAAAG | Rev PureA-GAATTCATG/ComFC |
| Op614 | GATTACAAGGATGACGACGATAAGTAAAGCGGCCGCGACTCT AGATCATAATCAGCC | For FLAG* - linker |
| Op853 | GAGAATATTGTAGGAGATCTTCTAGAAA**GAT**AAAGAGGGCTT AAAACAGCGCTTAAGCC | For Eco47- HP1473 |
| Op854 | CCGGATGGCTCGAGTTTTTCAGCAAGATTCAGGGCGGTTACCC CCTAAACC | Rev HP1473 dw-Eco47 |

| | | |
|---|---|---|
| Op855 | GGCTTAAGCGCTGTTTTAAGCCCTCTTTATCTTTCTAGAAGATC TCCTACAATATTCTC | Rev Eco47- HP1473up |
| Op856 | GGTTTAGGGGGTAACCGCCCTGAATCTTGCTGAAAAACTCGAG CCATCCGG | For HP1473dw -Eco47 |
| Op857 | TCGCCGCTTTTATAAAATGCGCTGTTTAACCTGTTTGGTACCCG GGTGACTAACT | For HP1473up /20nt HP1473-Apra |
| Op858 | TTAAAAAATAAAATTATAACTCATTCATCCGCGCTGCAAAGGAT CCCCGTGTCATTATT | Rev Apra- 20nt HP1473+HP1473dw |
| Op859 | AATAATGACACGGGGATCCTTTGCAGCGCGGATGAATGAGTT ATAATTTTATTTTTTAA | For Apra- 20nt HP1473+Hp1473dw |
| Op860 | AGTTAGTCACCCGGGTACCAAACAGGTTAAACAGCGCATTTTA TAAAAGCGGCGA | Rev HP1473up /20nt Hp1473-Apra |
| XV2 | TGGGTGAACCTGCAGGTGGGCAAAGATGTCCTAGCAATGTAA TCGTCAAGCTTTATGCCGTT | 62 mer ssDNA5-Cy5 labelled |
| cXV2 | AACGGCATAAAGCTTGACGATTACATTGCTAGGACATCTTTGC CCACCTGCAGGTTCACCCA | 62 mer ssDNA Complementary to XV2 |

**Table S4.**
**Plasmids used in this study**

| Name | Description | Source |
|---|---|---|
| p978 | pJet1.2-*hp1473::Cm* | This work |
| P1175 | pJet1.2-*RdxA:: Km* | Lab collection |
| p1176 | pJet1.2-*RdxA:: Prom-hp1473-Km* | This work |
| p1204 | pJet1.2-*RdxA:: Prom-hp1473-T165A-Km* | This work |
| P1284 | pJet1.2-*RdxA:: Prom-hp1473-FLAG-Km* | This work |
| P1310 | pJet1.2-*RdxA:: Prom-hp1473-C15SC18S-FLAG-Km* | This work |
| P1410 | pET21-*His6-TEV- hp1473* | This work |
| P1412 | pET21-*His6-TEV- hp1473-T165A* | This work |
| P1088 | pJet1.2-PromUreA-Cm | Lab collection |
| P1672 | pJet1.2-PromUreA-*hp1473*-Flag-Cm | This work |
| P1674 | pJet1.2-PromUreA-*hp1473-T165A*-Flag-Cm | This work |
| P1676 | pJet1.2-PromUreA-*hp1473-C15S C18SFlag*-Cm | This work |
| P1699 | pJET1.2-*hp1743::Apra* | This work |

**Supplementary movies 1-3.** Representative movies showing internalisation of the ATTO488-labelled transforming DNA (red) into the cytoplasm of GFP expressing *H. pylori* strains (green): *Wild-type* (**Supplementary movie 1)**, *ΔcomEC* (**Supplementary movie 2)**, and *ΔcomF* (**Supplementary movie 3)**. Sample preparation, image acquisition and processing were performed as described in Corbinais et al., 2016.
